# Force-regulated spontaneous conformational changes of integrins α_5_β_1_ and α_V_β_3_

**DOI:** 10.1101/2023.01.09.523308

**Authors:** Yunfeng Chen, Fang Kong, Zhenhai Li, Lining Arnold Ju, Cheng Zhu

## Abstract

Force can modulate the properties and functions of macromolecules by inducing conformational changes, such as coiling/uncoiling, zipping/unzipping, and folding/unfolding. Here we compared force-modulated bending/unbending of two purified integrin ectodomains, α_5_β_1_ and α_V_β_3_, using single-molecule approaches. Similar to previously characterized mechano-sensitive macromolecules, the conformation of α_5_β_1_ is determined by a threshold head-to-tail tension, suggesting a canonical energy landscape with a deep energy well that traps the integrin in the bent state until sufficient force tilts the energy landscape to accelerate transition to the extended state. By comparison, α_V_β_3_ exhibits bi-stability even without force and can spontaneously transition between the bent and extended conformations in a wide range of forces without energy supplies. Molecular dynamics simulations revealed consecutive formation and disruption of 7 hydrogen bonds during α_V_β_3_ bending and unbending, respectively. Accordingly, we constructed an energy landscape with hexa-stable intermediate states to break down the energy barrier separating the bent and extended states into smaller ones, making it possible for the thermal agitation energy to overcome them sequentially and to be accumulated and converted into mechanical work required for α_V_β_3_ to bend against force. Our study elucidates the different inner workings of α_5_β_1_ and α_V_β_3_ at the sub-molecular level, sheds lights on how their respectively functions are facilitated by their distinctive mechano-sensitivities, helps understand their signal initiation processes, and provides critical concepts and useful design principles for engineering of protein-based biomechanical nanomachines.

## Introduction

Integrins are a family of heterodimeric transmembrane molecules on the surface of nearly all cells. By mediating cell-cell and cell-matrix adhesion and bi-directional transmembrane signal transduction, integrins play key roles in many cellular functions, regulating cell attachment, migration, proliferation differentiation and more^1^. Dysregulation of integrins is often associated with diseases such as cancer, immune disorders, and vascular thrombosis^2^. Many cells use two key integrins – α_5_β_1_ and α_V_β_3_ – to bind the extracellular matrix (ECM) and form focal adhesion, but their functions are distinct: α_5_β_1_ molecules translocate laterally and cluster to support firm adhesion and cell spreading, whereas α_V_β_3_ molecules remain relatively stationary in focal adhesion and mediate early-stage mechanotransduction and rigidity sensing^3-5^. The molecular basis of such functional distinction is unclear, although it has been vaguely suggested to be associated with the structural differences in α_5_β_1_ and α_V_β_3_ ectodomains^4,5^.

We previously show that integrin α_L_β_2_ and α_V_β_3_ undergo force-modulated conformational changes, such that force facilitates unbending but suppressed bending, shifting the conformational equilibrium towards extension^6,7^. From a mechanical perspective, however, it is surprising that integrins can spontaneously bend against a wide range of forces. It is intuitive that a head-to-tail tension can induce a bent integrin to unbend regardless of which conformation is more stable prior to force application, because force can always tilt the energy landscape to force the extended conformation to become more stable, if this is not already the case in the absence of force. However, even if the bent conformation is more stable, thus the extended integrin tends to bend in the absence of force, its spontaneous bending against a tensile force is counterintuitive when the tension is as high as 10-20 pN. The mechanical work done by the integrin to decrease the head-to-tail distance from 20 to 13 nm under a linearly increasing force from 20 to 22.1 pN (for a force transducer spring constant of 0.3 pN/nm, see Methods) is ∼35 *k*_B_*T*, comparable to the free energy of biotin-avidin binding (∼35 *k*_B_*T*), one of the strongest noncovalent interactions^8^.

Our previous experiments on force-modulated integrin bending and unbending were performed on living cells^6,7^, where cell activity can regulate integrin conformational changes biologically. It seems natural to hypothesize that it must be the cell who provides an “active energy” to bend the integrin against force, since the energy from the environmental thermal agitation (0.5 *k*_B_*T* for each degree of freedom of the integrin motion) seems insufficient to provide 35 *k*_B_*T* energy. However, it is difficult to envision how this presumably cell-provided energy is converted into mechanical work to power the bending of the extended integrin, which occurs distal from cell surface.

To test the above hypothesis, we used two force spectroscopic techniques to perform single-molecule experiments on purified α_5_β_1_ and α_V_β_3_ proteins and demonstrated in cell-free systems that, while both integrins were able to undergo spontaneous bending and unbending under force without the supply of energy such as ATP, their conformational changes exhibited distinctive mechanical and kinetic properties, which may be related to their distinctive structures and biological functions^9^. Specifically, the α_5_β_1_ conformation was mostly bent in Ca^2+^ and mostly extended in Mn^2+^, suggesting a canonical energy landscape with a deep energy well that traps the integrin in the bent state in Ca^2+^ and the extended state Mn^2+^. Force could tilt the energy landscape to shift the system in Ca^2+^ to bi-stable such that α_5_β_1_ would spontaneously transition back-and-forth between the bent and extended states. In sharp contrast, α_V_β_3_ might take either bent or extended conformation in both Ca^2+^/Mg^2+^ and Mn^2+^, and the extended α_V_β_3_ could spontaneously bend back against a wide range of tensile forces in similar ways as cell surface α_V_β_36_, falsifying our “biological energy” hypothesis and suggesting a physical mechanism. To explain the unusual behaviors of α_V_β_3_, we developed a multi-state conformational energy landscape for this integrin, which was supported by molecular dynamics simulations and able to fit our experimental data well. Our work reveals that α_V_β_3_ protein can spontaneously bend and unbend in a wide range of mechanical forces in the absence of biochemical stimuli and biological energy, therefore providing critical concepts and useful design principles for engineering of protein biomechanical machines.

## Results

### Directly observing single α_5_β_1_ protein unbending and rebending

Using atomic force microscopy (AFM), we tested whether mechanical force can induce conformational changes of α_5_β_1_ protein independent of cell regulation. Recombinant α_5_β_1_ ectodomain with a human IgG Fc tag (α_5_β_1_-Fc) was captured on a polystyrene surface (Fig. 1A), and driven to touch the fibronectin module III domains 7–10 (FN, containing both the RGD sequence and synergistic site^10^) adsorbed on a cantilever tip to allow for bond formation (Methods). A tensile force was loaded on each α_5_β_1_–FN bond, which was ramped by retracting the polystyrene surface at 200 nm/s until reaching 20 pN, and then unloaded to 0 pN at the same rate (Fig. 1B). Inspection of the force vs time traces often reveals a clearly visible kink in the middle of both the loading and unloading phases, where the slope of the curve suddenly drops from positive to zero or even negative in the loading phase and abruptly jumps from negative to zero or even positive in the unloading phase (Fig. 1B). These kinks are clear indications of protein conformational changes such as unfolding-refolding^11^.

**Figure 1.**
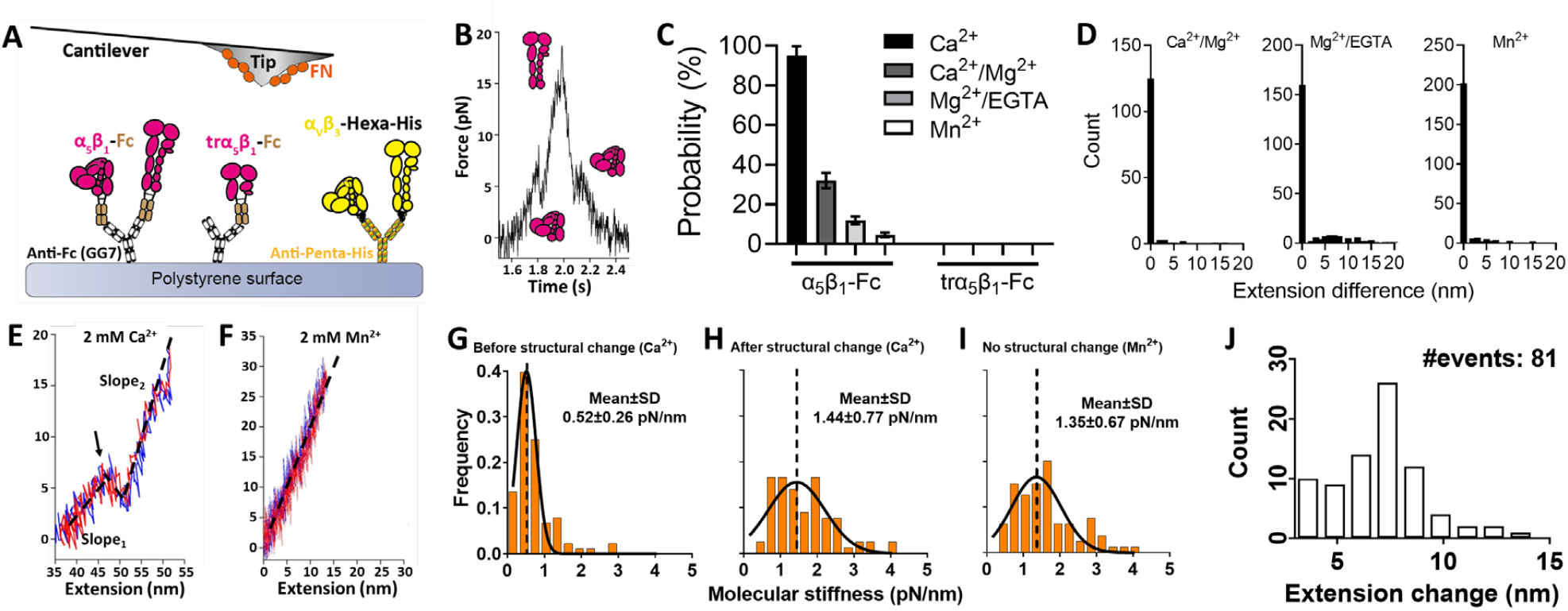
Observation and characterization of force-modulated α_5_β_1_ unbending and bending by AFM. A. Superimposition of AFM experimental setups for α_5_β_1_ and α_V_β_3_. Recombinant α_5_β_1_, truncated α_5_β_1_, and α_V_β_3_ were respectively immobilized on a polystyrene surface using capturing mAbs GG7 (anti-Fc) or anti-Hexa-Histidine. The polystyrene surface was brought in and out of contact with a cantilever coated with FN to allow integrin–ligand bond formation. Full-length α_5_β_1_ and α_V_β_3_ ectodomains, but not truncated α_5_β_1_, can adopt both bent and extended conformations. B. A representative force vs. time trace of loading-unloading cycle on a α_5_β_1_–FN bond. Two “kinks”, one in the loading and the other in the unloading phase, respectively repersent integrin unbending and bending along with cartoons depicting different α_5_β_1_ conformations prior and after the kinks. C. Mean ± standard error of the probability of observing structural changes in α_5_β_1_ or trα_5_β_1_ in a force loading-unloading process in different metal ion conditions. D. Distribution of the difference of the α_5_β_1_/FN complex molecular length before and after a full force loading-unloading cycle. E,F. Representative force vs. extension curves of loading (*red*) and unloading (*blue*) in Ca^2+^ (E) and Mn^2+^ (F), obtained by eliminating the time variable from the force vs. time trace and the corresponding piezo movement vs. time trace. “Kinks” (*arrow*) were much more frequently observed in Ca^2+^ than in Mn^2+^, as respectively exemplified in (E) and (F). The loading and unloading traces were linearly fitted (*black dashed lines*) to evaluate molecular stiffness, showing in (E) clearly stiffness increase from *Slope*_1_ to *Slope*_2_ when α_5_β_1_ changed from the bent to extended conformation in the loading phase, and stiffness decrease from *Slope*_2_ to *Slope*_1_ when α_5_β_1_ changed from the extended to bent conformation in the unloading phase. Neither kink nor slope change was observed in the londing and unloading phases in (F). G-I. Histograms of the α_5_β_1_–FN complex stiffness before (G) and after (H) unbending in Ca^2+^, and with no visible structural change in Mn^2+^ (I), and their respecetively Gaussian distribution fits (mean and standard deviation (SD) annotated). J. Histogram of the α_5_β_1_ head-to-tail molecular extension change due to unbending in Ca^2+^.

To identify the origin of these conformational changes, we first replaced the full-length α_5_β_1_-Fc with a truncated construct that contains only the α_5_β_1_ headpiece (trα_5_β_1_-Fc; Fig. 1A). In all four cation conditions – 2 mM Ca^2+^ (Ca^2+^), 1 mM Ca^2+^ plus 1 mM Mg^2+^ (Ca^2+^/Mg^2+^), 1 mM Mg^2+^ plus 1 mM EGTA (Mg^2+^/EGTA) and 2 mM Mn^2+^ (Mn^2+^) – that favor different global integrin conformations, the conformational changes seen with the full-length α_5_β_1_-Fc were no longer observed (Fig. 1C), indicating that these conformational changes are from the α_5_β_1_ ectodomain but not the recombinant Fc tail, GG7 or FN, and require the α_5_β_1_ tailpiece. Secondly, such conformational changes occurred progressively less frequently as the cation composition changed to those that activate integrins more and more potently, resulting in a frequency hierarchy of Ca^2+^ > Ca^2+^/Mg^2+^ > Mg^2+^/EGTA > Mn^2+^ (Fig. 1C). These results suggest the observed structural lengthening/shortening events to be those of integrin unbending/rebending, because this explains the frequency hierarchy: the activating cation conditions facilitate more integrins to adopt the extended conformation, leaving less integrins in the bent conformation capable of unbending. Thirdly, in all cation conditions, the structural lengthening in the loading phase was almost always ensued by a structural shortening in the unloading phase. By plotting force against the molecular extension of the bond, we found that the molecular complex fully recovers to its original length at the end of the loading-unloading cycle with no hysteresis (Fig. 1D) and the loading and unloading phases largely overlap (Fig. 1E), suggesting that the conformational changes are highly reversible and ruling out the alternative interpretation that they represent irreversible structure denaturation. Fourthly, the slope of the force-extension curve was seen to always increase after a structural lengthening and decrease after a structural shortening. Since such slope represents molecular stiffness (Methods), these findings indicate that α_5_β_1_ becomes stiffer after the structural lengthening and becomes softer after the structural shortening (Fig. 1F, H, I). This is consistent with our previous results that integrins are stiffer in their extended conformation than the bent conformation^6,7,12^. Using molecular stiffness as a signature readout of integrin conformation, we found that the value of the post-extension stiffness is similar to those integrins showing no structural changes in Mn^2+^ (Fig. 1G-I), which agrees with our hypothesis that integrins would be nearly unable to unbend or bend in this cation condition, because Mn^2+^ has already activated most of the integrins to the extended conformation. Finally, the molecular extension change due to structural lengthening centers around 7.5 nm (Fig. 1J), comparable to the head-to-tail length increase of an unbending integrin α_5_β_1_ characterized by theoretical modeling^13^. Together, these results indicate that the conformational changes observed in the force vs time and force vs extension curves originate from integrin α_5_β_1_ unbending and rebending.

### Directly observing single α_V_β_3_ protein bending and unbending

The biophysical characteristics of force-modulated α_5_β_1_ unbending and rebending are very different from those previously characterized for integrins α_L_β_2_ and α_V_β_3_ on the cell surface. The force range within which unbending and rebending events could be observed is quite narrow for α_5_β_1_ (<10 pN, Fig. 1C, F; also see Fig. 5A-D below) but much wider for α_L_β_2_ and α_V_β_3_ (up to 40 pN)^6,7^. The range of head-to-tail extension changes generated by unbending and bending was also much smaller for α_5_β_1_ (3-14 nm, Fig. 1J) than α_L_β_2_ and α_V_β_3_ (5-30 nm)^6,7^. Furthermore, the kinetics of conformational changes of α_5_β_1_ protein were also very different from those of cell surface α_L_β_2_ and α_V_β_3_. (Un)bending kinetics of the α_5_β_1_ protein will be characterized and compared to those of the α_V_β_3_ protein later, but their quantification has been described previously by two pairs of parameters^6,7^. Time-to-switch *t*_0±_ is the waiting time required for conformational switch to occur. Switching time *t*_sw±_ is the time taken for the conformational switch from start to finish. Using these definitions, we found that the kinetics are much more rapid for α_5_β_1_ (e.g., *t*_0-_ at 5 pN is ∼0.02 s) than α_L_β_2_ and α_V_β_3_ (up to 3 s)^6,7^. A hypothetical explanation for the different (un)bending behaviors observed here and those previously quantified might be the absence of cell regulation for the α_5_β_1_ ectodomain but the presence of cell regulation for the full-length α_L_β_2_ and α_V_β_3_. To test this hypothesis, we applied the same AFM approach to study the conformational changes of the α_V_β_3_ ectodomain bound to FN. Surprisingly, no “kink”, i.e., sudden slope changes in the force-time curves was observed over hundreds of force loading-unloading events; clamping the α_V_β_3_–FN bonds under a constant force or applying cyclic forces did not yield any kink type of conformational changes either.

Cell surface α_V_β_3_ (un)bending events were previously observed using another force spectroscopy technique: the biomembrane force probe (BFP, Fig. 2A)^6^. We reasoned that the observation of α_V_β_3_ conformational changes might not be favored by the stiff AFM cantilever but favored by the soft BFP force sensor (spring constants ∼3 vs ∼0.3 pN/nm), because a 10 nm head-to-tail length change caused by α_V_β_3_ bending would result in a ∼30 pN force increase in the AFM, which would severely inhibit bending, but only ∼3 pN force change in the BFP, which would not. We thus used the BFP, consisting of a micropipette-aspirated human red blood cell (RBC) with a streptavidin (SA) and FN co-functionalized probe bead attached to its apex to serve as an ultrasensitive force transducer (Fig. 2A *left*). A bead coated with recombinant α_V_β_3_ protein was aspirated by an opposing micropipette (Fig. 2A, *right*) and driven to repeatedly contact the probe bead to induce binding (Methods). Binding was abolished by an α_V_β_3_-blocking mAb, LM609, confirming the specificity of α_V_β_3_–FN interaction (Fig. 2B).

**Figure 2.**
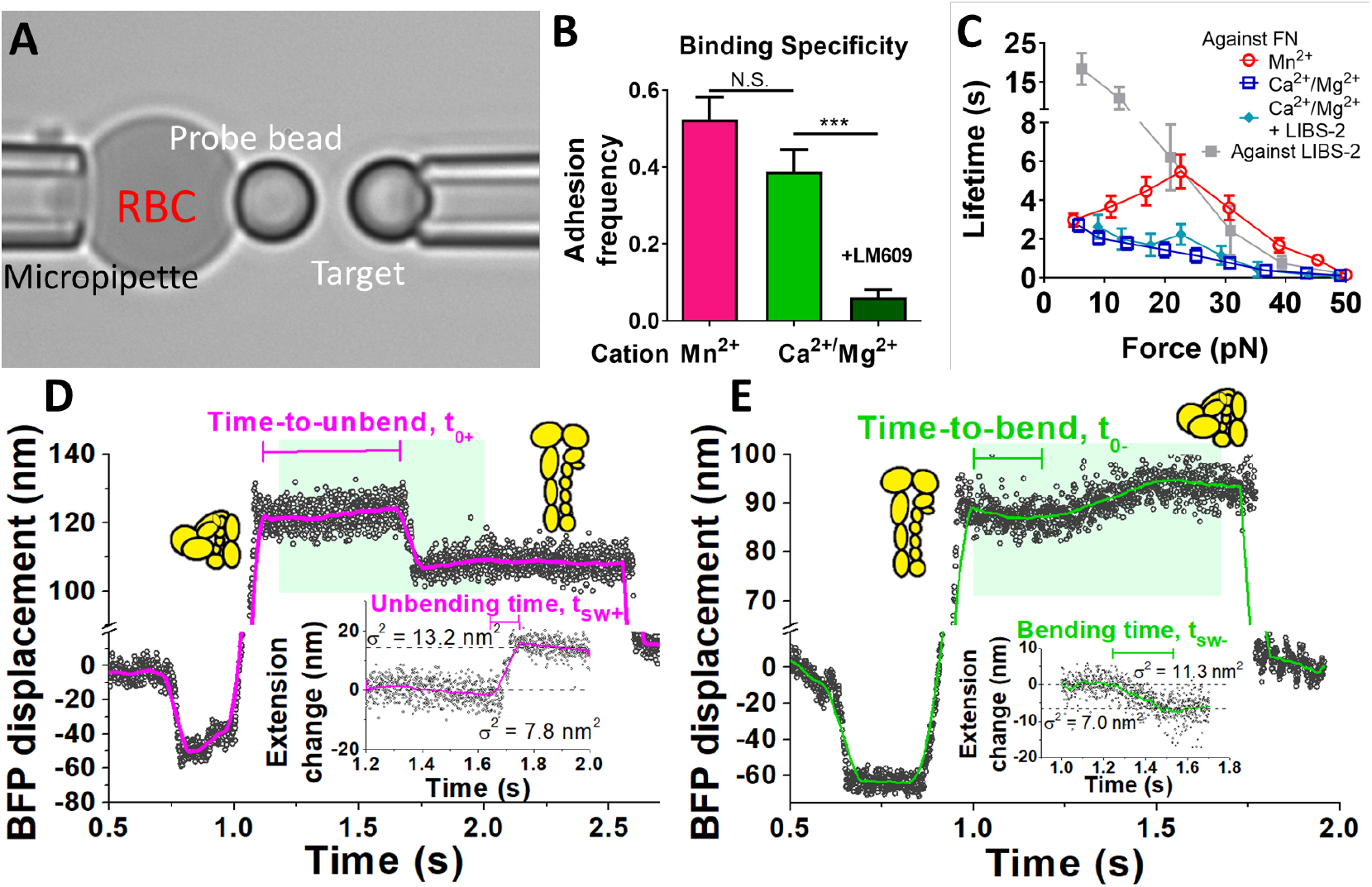
Observation of force-regulated α_V_β_3_ unbending and bending by BFP. A. BFP photomicrograph. A force probe was composed of a micropipette-aspirated biotinylated RBC and a SA and FN co-coated glass bead attached to its apex. A bead coated with α_V_β_3_ ectodomain was aspirated by an opposing micropipette as the target. B. Adhesion frequency of α_V_β_3_–FN binding in Mn^2+^ and Ca^2+^/Mg^2+^. The addition of blocking mAb LM609 blocked most of the adhesion events in Ca^2+^/Mg^2+^. C. Mean ± s.e.m. of lifetime vs. force of single α_V_β_3_–FN bonds in indicated conditions or α_V_β_3_–LIBS-2 bonds. D,E. Representative BFP displacement vs. time trace showing an integrin unbending (D) and bending (E) event in the position-clamp phase, along with cartoons depicting different α_V_β_3_ conformations before and after (un)bending. The data (*points*) is smoothened using the Savitzky-Golay method (*curves*) to obtain a higher force resolution. Inserts: Detailed views of the conformational changes within the cyan-shaded windows that convert the BFP displacement to the α_V_β_3_ extension change, which include the standard deviations of the signals, σ, as a measure of thermal fluctuation before and after the (un)bending. Definition of time-to-switch and switching time are indicated.

After titrating the FN coating on probe beads to lower the adhesion frequency to 20%, a necessary condition for most binding events to be mediated by single bonds^14^, α_V_β_3_ was then interrogated under both Ca^2+^/Mg^2+^ and Mn^2+^ conditions using distance-clamping assay^6^: the α_V_β_3_–FN bond was first pulled to a certain force level, and target bead was then clamped at a fixed position. The bonds could sustain a wide range of forces, with lifetimes much longer in Mn^2+^ than in Ca^2+^/Mg^2+^, consistent with the activating role of Mn^2+^ (Fig. 2C). Unlike purified α_5_β_1_–FN interaction that forms catch-slip bonds not only in Mn^2+^, but also in Ca^2+^/Mg^2+^ and Mg^2+^/EGTA^15^, purified α_V_β_3_–FN interaction only formed a catch-slip bond in Mn^2+^, but a slip-only bond in Ca^2+^/Mg^2+^. This slip-only bond indicates limited effect of sustained force to strengthen α_V_β_3_ bonding to FN, which agrees with our previously reported weak α_V_β_3_–FN catch-slip bond on the cell surface^6^.

In the clamping phase of some lifetime measurements, we observed α_V_β_3_ unbending or bending events, respectively signified by a concurrent decrease in the mean force and increase in the force fluctuation or a concurrent increase in the mean force and decrease in the force fluctuation^6,7^(Fig. 2D,E; statistics summarized in Supp. Table 1). Unlike purified α_5_β_1_ and consistent with cell surface α_V_β_3_, the conformational changes of purified α_V_β_3_ occurred under a wide range of forces (Fig. 3A,B) with relatively long time-to-switch (*t*_0+_ and *t*_0–_ respectively for unbending and bending) and switching time (*t*_sw+_ and *t*_sw–_ respectively for unbending and bending)^6^ (c.f., Fig. 2D,E). Replacing FN on the probe beads with LIBS-2, a mAb that binds the α_V_β_3_ βTD domain at its tailpiece^16^, abolished the above signature signals for integrin conformational changes (Supp. Table 1) despite the long lifetimes (Fig. 2C), ruling out the alternative possibility that the putative bending/unbending events are due to multiple bond rupture/formation or instrument drifting. Adding LIBS-2 to the solution, which stabilizes β_3_ integrins in the extended conformation^17^, also eliminated all bending events (Supp. Table 1). Interestingly, LIBS-2 did not significantly alter the α_V_β_3_–FN bond type and lifetimes in Ca^2+^/Mg^2+^ (Fig. 2C), decoupling integrin extension and catch-slip bond formation. During (un)bending, the change in the RBC elongation (Fig. 2D,E, *cyan shaded areas*) is equal to the change in the integrin head-to-tail length^6,7^. These length changes of both unbending and bending events follow a single-Gaussian distribution (Fig. 3A,B) with an indistinguishable average value of ∼13 nm in both Ca^2+^/Mg^2+^ and Mn^2+^ (Fig. 3C), again agreeing with our previous observations on cell surface α_V_β_3_^6^.

**Figure 3.**
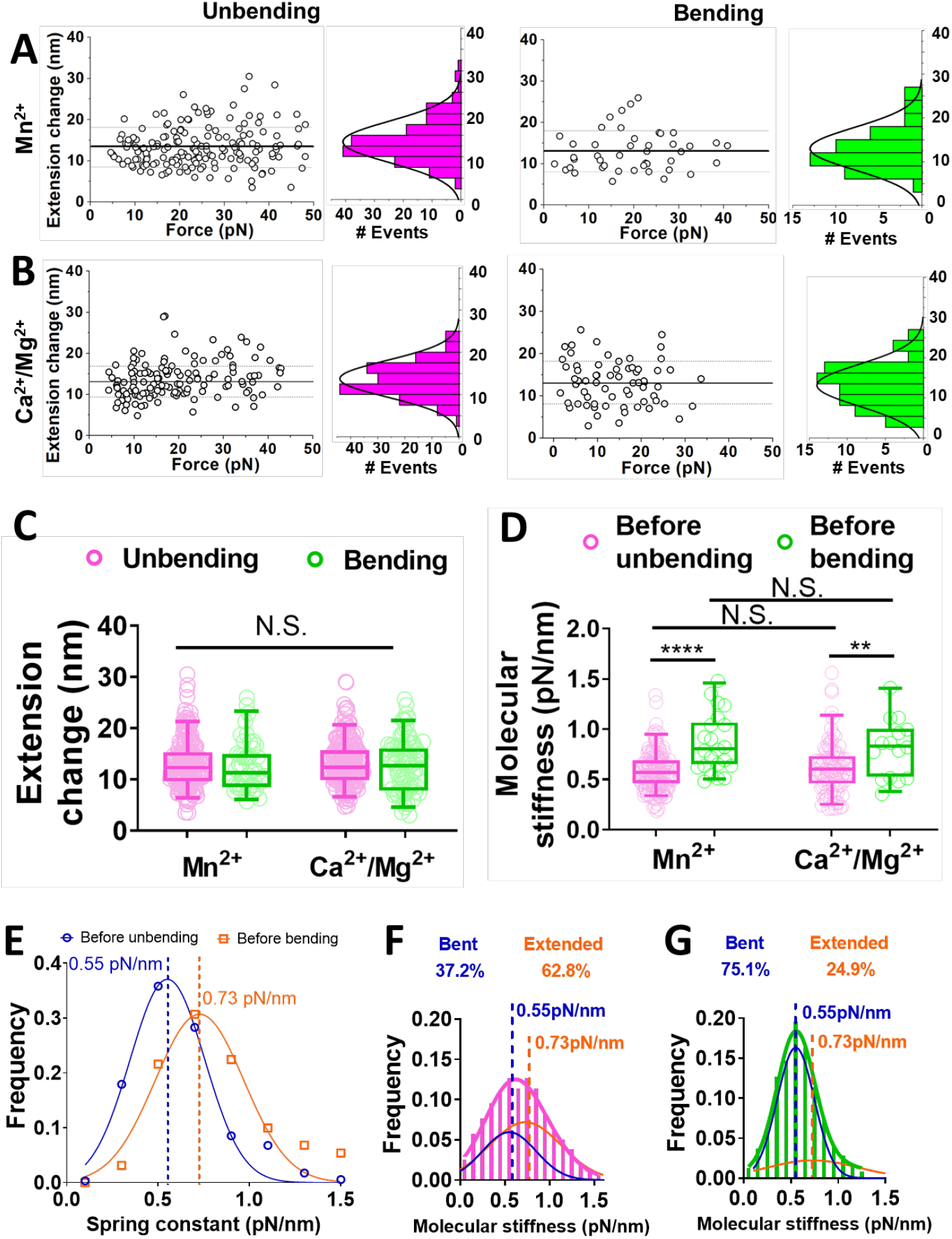
Characterization of force-regulated α_V_β_3_ unbending and bending by BFP. A,B. Scatter plots and Histograms (*bars*) and Gaussian fits (*curves*) of α_V_β_3_ extension changes due to unbending (*left*) and bending (*right*) in Mn^2+^ (A) and Ca^2+^/Mg^2+^ (B). C. Data (*points*) and the median and 5-95 percentiles (box and whisker) of α_V_β_3_ extension changes due to unbending and bending in Mn^2+^ and Ca^2+^/Mg^2+^. D. Data (*points*) and the median and 5-95 percentiles (box and whisker) of the α_V_β_3_–FN molecular stiffness before unbending events and before bending events. N.S. = not significant assessed by one-way ANOVA; ** *p* <0.01; **** *p* <0.0001, assessed by unpaired, two-tailed Student’s t-test. E. Fitting the α_V_β_3_/FN molecular stiffness before unbending and before bending with Gaussian distribution to respectively acquire the average molecular stiffness associated with bent and extended integrins. F,G. Fitting the α_V_β_3_/FN molecular stiffness in Mn^2+^ (F) and Ca^2+^/Mg^2+^ (G) with dual-Gaussian distribution to calculate the proportions of BFP-detected integrins in bent and extended conformations. The means of the two Gaussian distributions, respectively associated with bent and extended α_V_β_3_, were derived from (E).

Furthermore, the stiffness of the α_V_β_3_–FN complex is lower before unbending than before bending (Fig. 3D), consistent with the signature integrin stiffening upon unbending^6,7,12^. Since the stiffness depends only on the conformation but not cation condition (Fig. 3D), we pool data from both cation conditions together to examine the stiffness distributions for the bent and extended integrins, finding their respective means and standard deviations of 0.55 ± 0.20 and 0.73 ± 0.24 pN/nm (Fig. 3E), comparable to the values previously measured from cell surface α_V_β_3_^6^. Moreover, we plotted the histograms of additional stiffness measurements from each cation condition, regardless of whether integrin (un)bending events were observable, and fitted each by a dual-Gaussian distribution using 0.55 and 0.73 pN/nm as the two means to calculate the proportions of integrins in the bent and extended states. We found that, of those α_V_β_3_ that formed bonds, 62.8% were in the extended conformation in Mn^2+^, but 75.1% were in the bent conformation in Ca^2+^/Mg^2+^ (Fig. 3F,G), consistent with the activating role of Mn^2+^. More importantly, these results confirm that integrin α_V_β_3_, unlike α_5_β_1_, is already bi-stable under zero force. Further, the data confirm that purified integrin α_V_β_3_ protein can spontaneously transition between the bent and extended conformations under a wide range of tensile forces in the absence of cellular regulation and biological energy supply.

### Integrin α_V_β_3_ showed no CMR effect

In our position-clamp experiment, spontaneous unbending and bending of α_V_β_3_ respectively decreased and increased the force on its bonds with FN (Fig. 2D,E), which may help strengthen the bonds through a mechanism called “cyclic mechanical reinforcement” (CMR) where a cyclic force applied to a receptor–ligand bond greatly prolongs its lifetime. CMR was initially observed with integrin α_5_β_1_–FN bonds^18^, but later also observed with actin–actin bonds^19^. To test the CMR effect on α_V_β_3_–FN bonds, we first used AFM (Fig. 1A, Methods) as did previously on α_5_β_1_–FN bonds^18^. Once a bond was detected, two types of cyclic forces were applied: 1) one loading-unloading cycle that first peaks at 20 pN and then drops to and holds at 5 pN (Fig. 4A); and 2) cyclic forces with zero, one, two or three complete loading-unloading cycles followed by ramping to and clamping at a peak force of 10 pN (Fig. 4C). Unexpectedly, neither type of cyclic forces prolonged α_V_β_3_–FN lifetimes (Fig. 4B,D), showing the lack of CMR effect.

**Figure 4.**
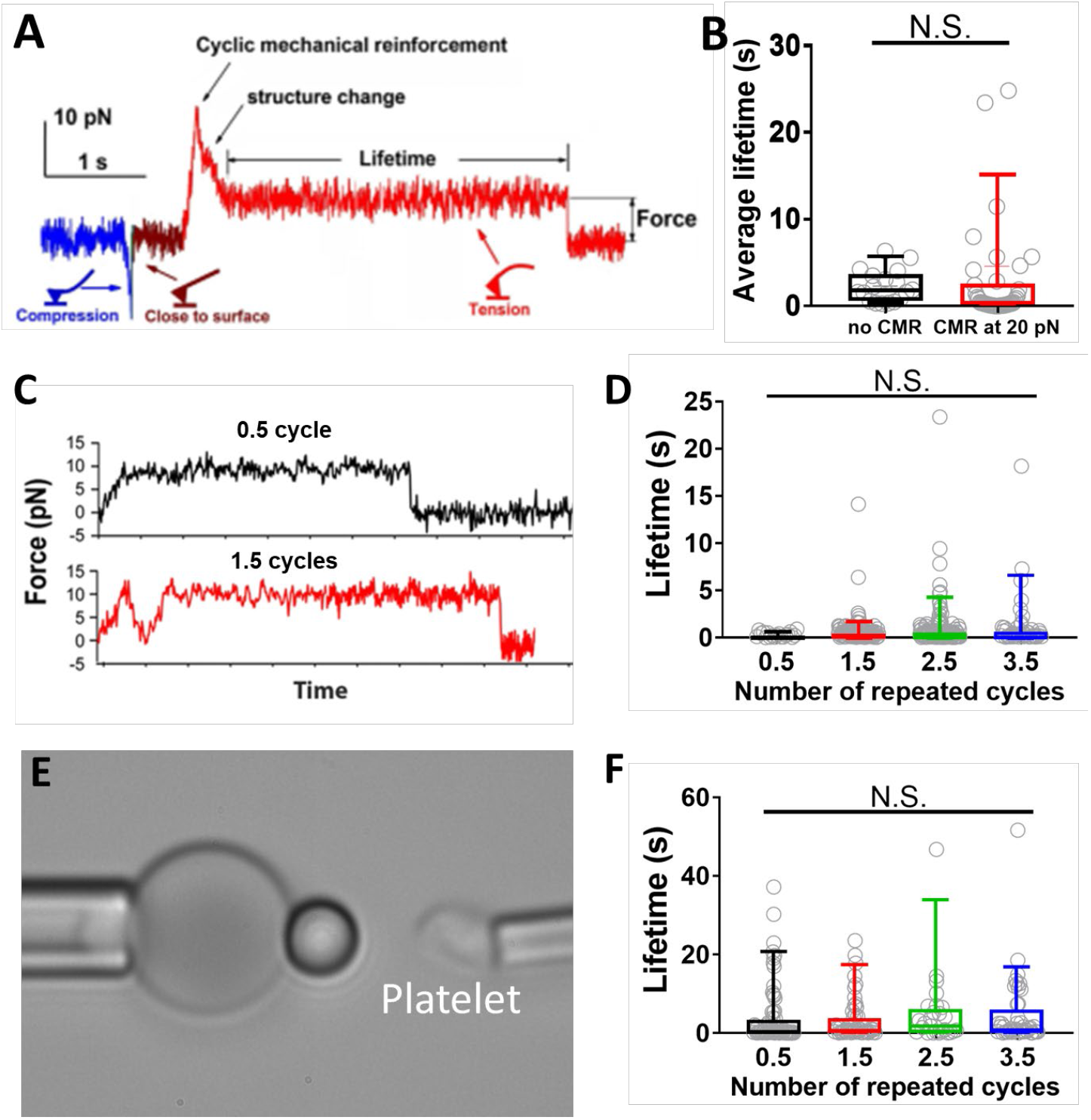
Measuring cyclic mechanical reinforcement (CMR) of integrin α_V_β_3_ using AFM and BFP. In both systems the ligand coating was titrated to reach infrequent adhesion (∼20%), a necessary condition for most adhesion events to be mediated by single-bonds. A. A representative AFM force vs. time trace showing a CMR with one loading-unloading cycle with a ∼20 pN peak force followed by bond lifetime measurement at ∼5 pN, which was used to generate the data in right group in (B) The cartoons indicated how the cantilever would be bent in different segments of the data curve. B. Data (*points*) and the median and 5-95 percentiles (box and whisker) of α_V_β_3_–FN bond lifetimes measured after a single-cycle CMR (*red*, exemplified in A) or without CMR (*black*, exemplified in C, *upper panel*). C. Two representative AFM force vs. time traces showing α_V_β_3_ lifetime measurements of a bond with 0.5 (*upper*) and 1.5 (*lower*) loading-unloading cycle before clamping at the peak force, which were used to generate the data in the first two groups in (D). D. Data (*points*) and the median and 5-95 percentiles (box and whisker) of α_V_β_3_–FN bond lifetimes measured after the indicated numbers of CMR cycles. E. BFP photomicrograph showing the experiment setup used to generate the data in (F), where a platelet aspirated by an opposing micropipette acted as the target. F. Data (*points*) and the median and 5-95 percentiles (box and whisker) of platelet α_V_β_3_–FN bond lifetimes measured after the indicated numbers of CMR cycles using the BFP showing in (E).

We also repeated the above experiments using BFP with α_V_β_3_-expressing platelets as the target (Fig. 4E). Inhibitory mAbs 10E5 and P1D6 were added to respectively block α_IIb_β_3_ and α_5_β_1_, two other FN-binding integrins on platelets, to ensure specific α_V_β_3_–FN interaction^12^. The second type of force loading-unloading cycles was applied to α_V_β_3_–FN bonds followed by ramping to and clamping at the peak force (10 pN). Despite the use of platelet α_V_β_3_, its bond lifetime with FN was still not prolonged by cyclic forces (Fig. 4F).

### Distinctive force-dependent kinetics of α_5_β_1_ and α_V_β_3_ conformational changes

We next analyzed the bending and unbending kinetics of α_5_β_1_ and α_V_β_3_, as characterized by the time-to-switch *t*_0±_ and switching time *t*_sw±_. In the first experiment, we employed AFM to pull α_5_β_1_ slowly (∼1 nm/s) after performing CMR to strengthen its bond with FN, which prolonged time for observation of repetitive unbending-rebending cycles in a single binding event^18^ (Fig. 5A), allowing us to collect ensembles of *t*_0±_ for kinetic analysis. Such consecutive back-and-forth events were indeed observed, but occurred at comparable frequencies only in a narrow force range (6-8 pN, Fig. 5A), as the probability or fraction of time during which the integrin stayed in the extended state increased rapidly and approached 1 as force exceeded 10 pN (Fig. 5B). This probability can be expressed as *P* = <*t*_0-_>/(<*t*_0+_> + <*t*_0-_>) because <*t*_0+_> and <*t*_0-_> are the respective average times required for the integrin to wait in the bent and extended states, respectively, until unbending and bending occurs, respectively. Also, the frequencies or kinetic rates of unbending and bending can be expressed as 1/<*t*_0+_> and 1/<*t*_0-_>, respectively. As expected, <*t*_0+_> decreased exponentially, and <*t*_0-_> increased exponentially, with increasing force *f* (Fig. 5C), behaving as a typical slip bond and catch bond, respectively^20^. We fitted the *P* vs *f* data to an existing two-state model^21,22^:

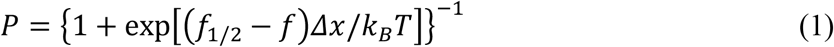

where *f*_1/2_ is the force at which the integrin has a 50-50 chance of staying in either the bent or extended state, Δ*x* is the separation distance between the two energy wells that define the bent and extended states in the energy landscape, and *k*_B_ is the Boltzmann constant, and *T* is absolute temperature. Fitting resulted in excellent agreement between the model and data (Fig. 5B), and returned *f*_1/2_ = 6.04 ± 0.01 pN, Δ*x* = 4.40 ± 0.06 nm, and *f*_1/2_*Δx*= 6.3 ± 0.09 *k*_B_*T*.

Eq. 1 can be derived by assuming the force-dependent <*t*_0+_> and <*t*_0-_> to follow the Bell model^23^ (Eq. 2a) and its “catch bond counterpart” (Eq. 2b):

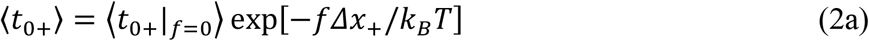

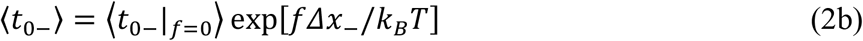

where ⟨ *t*_0±|*f*=0_ ⟩ are the respective values of ⟨ *t*_0±_ ⟩ at zero force, Δ*x*_±_ represent, respectively, the distances from the top of the energy barrier to the bottoms of the respective energy wells of the bent (Δ*x*_+_) and extended (Δ*x*_-_) conformations in the energy landscape at zero force. Substituting Eqs. 2a and 2b to the definition of *P* in terms of ⟨ *t*_0±_ ⟩ and comparing the result with Eq. 1, we obtain *Δx*= *Δx*_+_ + *Δ*_−_ and *f*_1/2_*Δx*= *k*_B_*T*In(⟨*t*_0+_|_*f*=0_⟩/⟨*t*_0−_|_*f*=0_⟩).

**Figure 5.**
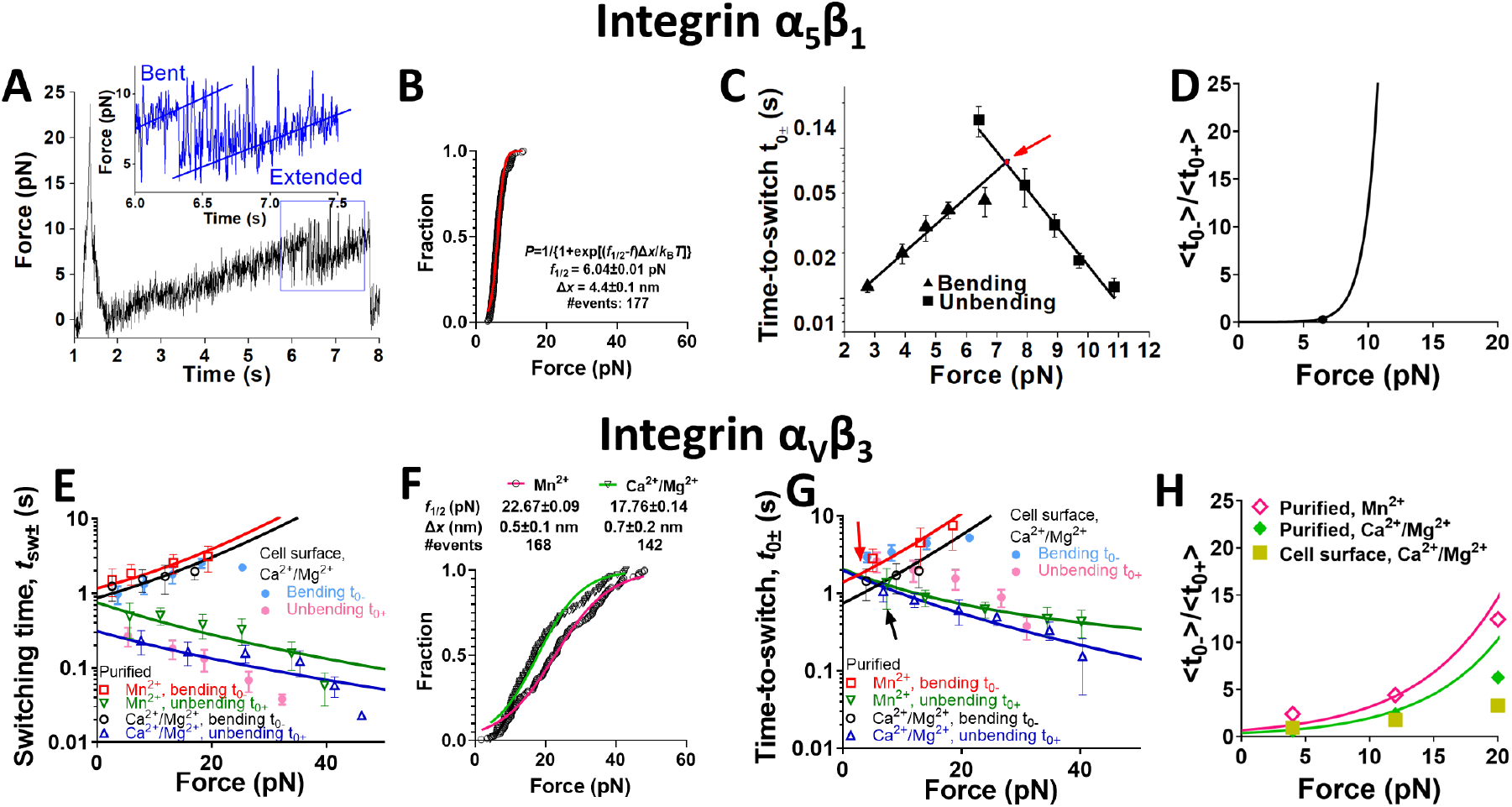
Force-modulated α_5_β_1_ and α_V_β_3_ bending and unbending kinetics. A. A representative force vs. time trace of applying slow ramping force on an α_5_β_1_–FN bond after a single cycle CMR, which was measured by AFM in Ca^2+^ to exemplify reversible and consecutive unbending-bending events of α_5_β_1_. Insert: zoom-in of the curve segment showing repeated bending-unbending events in a narrow force range near ∼7 pN. B. Cumulative histogram of integrin α_5_β_1_ unbending force distribution. The distribution was fitted by a theoretical model to derive the parameters of the energy landscape. The equation of the model and the derived parameters were denoted. C. Semi-log plots of mean ± s.e.m. α_5_β_1_ time-to-unbending *t*_0+_ (*square*) and time-to-bending *t*_0-_ (*triangle*) vs. force data and their fits by the Bell model (*curves*). The two fitting curves intersect at 7.4±0.6 pN and 0.076±0.017 s (*arrow*). D. Plot of <*t*_0-_> to <*t*_0+_> ratio of α_5_β_1_ ectodomain conformational changes, which represents the “equilibrium constant” between the extended and bent conformations vs. force, calculated based on experimental data (*point*) and the model fit in panel (C) (*curve*). E,G. Semi-log plots of mean ± s.e.m. α_V_β_3_ ectodomain unbending time *t*_sw+_ (E) or time-to-unbending *t*_0+_ (G) (*hollow triangle* and *hollow inverted triangle*) and bending time *t*_sw-_ (E) or time-to-bending *t*_0-_ (G) (*hollow square* and *hollow circle*) vs. force data measured in the indicated cation conditions, and their theoretical fits by the multi-state model described in the text. The *R*^2^ values of the fits are 0.95 and 0.96 for Mn^2+^ and Ca^2+^/Mg^2+^ conditions, respectively. Solid dots: mean ± s.e.m. *t*_0±_ and *t*_sw±_ vs. force of cell surface α_V_β_3_ unbending (*light magenta*) and bending (*light cyan*) events in Ca^2+^/Mg^2+^. F. Cumulative histogram of integrin α_V_β_3_ unbending force distribution with theoretical model fitting. H. Plots of <*t*_0-_> to <*t*_0+_> ratio of purified and cell surface α_V_β_3_ vs. force measured under indicated cation conditions and their model fits.

Directly fitting Eqs. 2a and 2b to the respective time-to-unbending and time-to-bending data in Fig. 5C yielded excellent agreement and returned ⟨*t*_0+_|_*f*=0_⟩ = 5.5 ± 2.9 s, Δ*x*_+_ *=* 1.6 ± 0.16 nm, ⟨*t*_0−_|_*f*=0_⟩ = 0.004 ± 0.0007 s, and Δ*x*_-_ *=* 2.4 ± 0.25 nm. The transition reached a balance at *f*_1/2_ = 7.4 ± 0.6 pN where times to both unbending 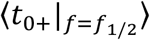 and bending 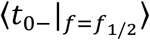 equaled 0.076 s. This *f*_1/2_ value, and the values of Δ*x*_+_ + Δ*x*_-_ = 4.0 ± 0.3 nm and *k*_B_*T* In(⟨*t*_0+_|_*f*=0_⟩/⟨*t*_0−_|_*f*=0_⟩) = 7.2 ± 0.56 *k*_B_*T* are comparable to those obtained by fitting Eq. 1 to data in Fig. 5B, supporting the quality of our data, the appropriateness of our model, and the reliability of the model parameters.

Across the *f*_1/2_ threshold, force quickly transitioned the integrin from the bent to extended conformation: as force increased from 4.3 to 10.5 pN, the dominant (>95%) integrin population rapidly changed from bent to extended conformation, which increased the population ratio of extended over bent integrins ⟨*t*_0−_⟩/⟨*t*_0+_⟩ by 400 folds, corresponding to an average force sensitivity of >60 folds/pN (Fig. 5D). Such a high force-sensitivity agrees with a previous theoretical study inferring that integrin α_5_β_1_ unbending is ultrasensitive to force^13^, reflecting nearly “digital” modulation of force on α_5_β_1_ conformation.

The kinetics of α_V_β_3_ conformational changes were then analyzed, which occurred over a much wider range of forces (Fig.5E-G). Compared to the Ca^2+^/Mg^2+^ cation condition, activating the integrin with Mn^2+^ resulted in slightly shorter ⟨*t*_sw+_⟩ and ⟨*t*_0+_⟩ as well as longer ⟨*t*_sw−_⟩ and ⟨*t*_0−_⟩ (Fig. 5E,G), consistent with the known coupling between unbending and integrin activation^24^. Comparing the transition dynamics of α_V_β_3_ and α_5_β_1_ revealed a sharp difference: while the conformational changes of α_5_β_1_ mimic a digital on/off switch (*t*_sw±_ always beyond the temporal resolution of 1 ms of our AFM instrument), the *t*_sw±_ of α_V_β_3_ was much longer and clearly force dependent: force prolonged ⟨*t*_sw−_⟩ but shortened ⟨*t*_sw+_⟩ (Fig. 5E). On the other hand, like α_5_β_1_, increasing force decreased ⟨*t*_0+_⟩ but increased ⟨*t*_0−_⟩ of α_V_β_3_ (Fig. 5G). Quantitatively however, the response of kinetics to force was much more moderate: within the force range of 4.0-28.0 pN where sufficient events were collected for statistical analysis, the population ratio of extended over bent α_V_β_3_ ⟨*t*_0−_⟩/⟨*t*_0+_⟩ only increased by 8.8 folds (a force sensitivity of ∼0.37 fold/pN) in Mn^2+^ and by 13 folds (a force sensitivity of ∼0.54 fold/pN) in Ca^2+^/Mg^2+^ (Fig. 5H), revealing a >100-fold greater resistance to force modulation than α_5_β_1_. We also re-analyzed our previously published data for cell surface α_V_β_3_^6^, finding results similar to cell-free α_V_β_3_ (Fig. 5E,G,H), indicating that these conformational changes are mainly modulated by force. Together, these results demonstrated distinctive mechanisms of force modulation on α_5_β_1_ and α_V_β_3_ (un)bending: a “digital” modulation for α_5_β_1_ and an “analogous” modulation for α_V_β_3_.

### Distinctive energy landscapes and transition modes of α_5_β_1_ and α_V_β_3_ conformational changes

The energy landscape for α_5_β_1_ unbending and bending, constructed based on the experimentally determined and model-fit parameters is depicted in Fig. 6A. Note that a force of *f*_1/2_ tilts the energy landscape such that the energy difference between the bent and extended states vanishes. As such, 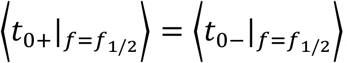, where α_5_β_1_ switches back and forth between the bent and extended conformation indefinitely using the energy from thermal agitations to hop over the force-suppressed energy barrier between the two states (Fig. 6A).

**Figure 6.**
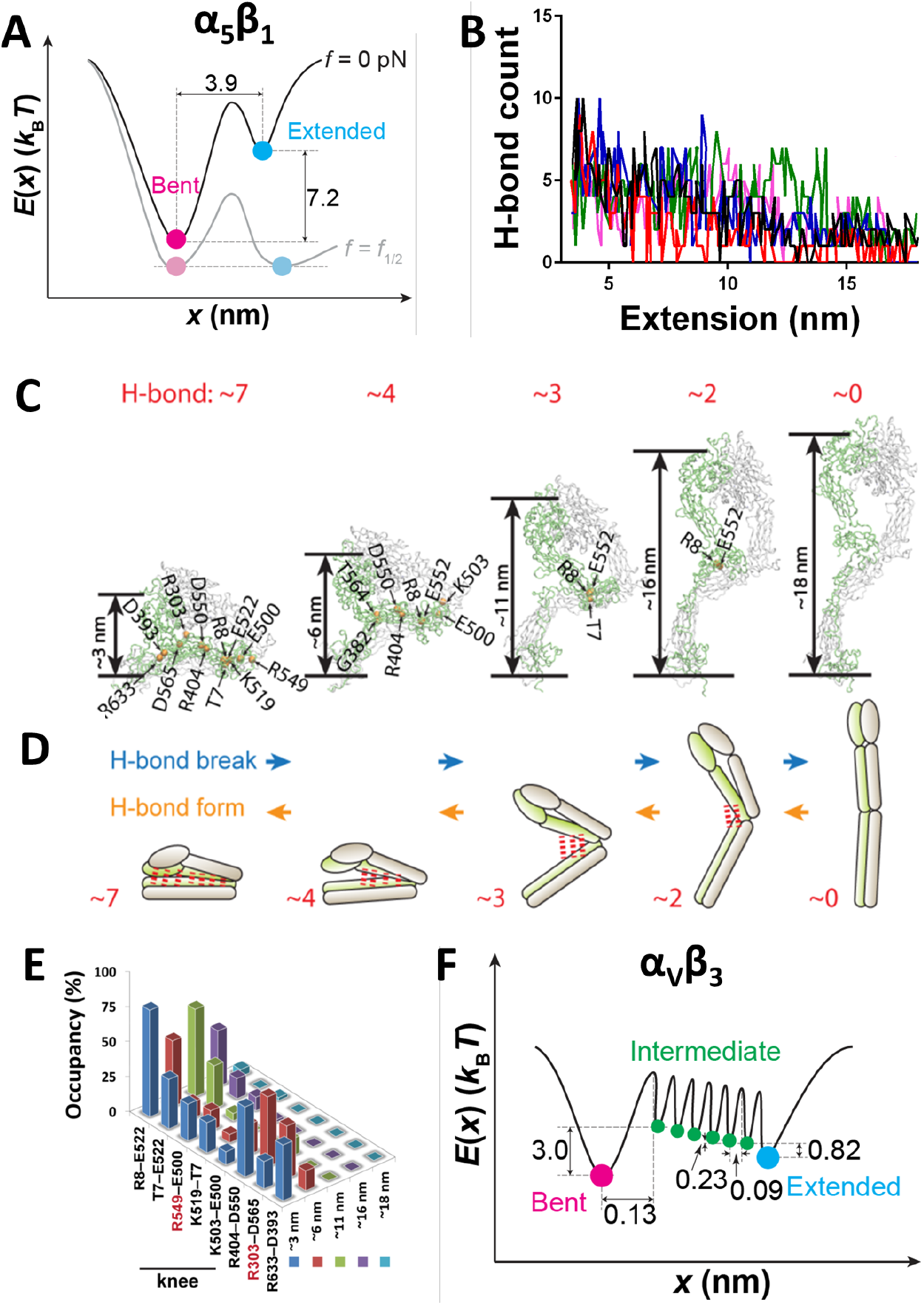
SMD simulations of α_V_β_3_ conformational changes and energy landscapes of α_5_β_1_ and α_V_β_3_. A. Energy landscapes of α_5_β_1_ ectodomain bending and unbending conformational changes under zero (*dark curve*) and *f*_1/2_ (*light curve*) force based based on the experimental and model-fit parameters (Fig. 5B). B. H-bond count of α_V_β_3_ during unbending collected from 5 runs of independent SMD simulations as indicated by different colors. C. Snapshots of representative α_V_β_3_ conformations (bent, three intermediate conformations, and extended) observed from the SMD simulations in which the most commonly observed H-bonds are indicated by their donor and acceptor residues. The red number on top of each panel represents the average H-bond number observed in 5∼6 independent MD simulations. D. Cartoons depicting the average numbers and locations of H-bonds in relation to the head-to-tail distances during α_V_β_3_ unbending. As the H-bonds are progressively disrupted, the integrin transitions from the bent to extended conformation (blue); as the H-bonds are progressively formed, the integrin transitions from the extended to bent conformation (*orange*). E. The average occupancy of the most frequently formed 8 H-bonds with α_V_β_3_ unbending to certain head-to-tail distances. Amino acids in α and β chains were shown in red and black, respectively. F. Energy landscapes of α_v_β_3_ bending and unbending in Mn^2+^ under zero force based on the experimental and model-fit parameters. Energy wells corresponding to the bent (*magenta*), extended (*cyan*) and intermediate (*green*) states are marked by different colors.

When constructing the energy landscape of α_V_β_3_ conformational changes, it was noted that the cumulative probability of its unbending against force *f* (Fig. 5F) can still be nicely fitted by Bell model^23^. However, observations that α_V_β_3_ is able to bend and unbend over a wide range of forces, including zero force, and that the switching times *t*_sw±_ are much longer than those taken by typical protein conformational changes (e.g., α_5_β_1_ bending/unbending observed here and talin and glycoprotein Ibα unfolding/refolding) and by conventional biochemical reactions^25^, indicate deviation from the conventional Bell model. We hypothesize that α_V_β_3_ conformational changes may involve a random sequence of formation and disruption of hydrogen bonds (H-bonds)^26,27^, resulting in a slow and complex sub-molecular process. To test this hypothesis, we conducted steered molecular dynamics (SMD) simulations to apply external forces to the α_V_β_3_ headpiece and observed that force would induce the gradual disruption of H-bonds originally present in the crystal structure (PDB code 3IJE) between the α_V_β_3_ headpiece and tailpiece during the unbending process (Fig. 6B). By holding the integrin at different head-to-tail distances against force and analyzing the disruption and formation of H-bonds over time, we found that in a bent α_V_β_3_, the headpiece could form as many as 134 H-bonds with the tailpiece, although most of them were formed with very low probabilities. The average number of H-bonds concurrently existing was 7, which gradually decayed as the integrin head-tail distance increased and reached 0 in the fully extended conformation (Fig. 6C,D and Supp. Fig. 1). Of the most frequently formed 8 H-bonds, 6 were within the integrin β subunit whereas the other 2 were between the α and β subunits. Five of these H-bonds contained a residue located in integrin knee region (Fig. 6E), suggesting the importance of the β integrin knee in the conformational change^28^. Notably, the H-bond most proximal to the integrin knee region (R8–E522) was not disrupted until the integrin reached full extension and was formed as soon as the integrin started to bend; by comparison, H-bonds distal to the knee region (e.g., R633-D393) were disrupted as soon as the integrin started to unbend, and only formed when the integrin was nearly fully bent (Fig. 6E). These results indicate a correlation between the H-bonds’ distance to the knee and the time sequence of their disruption and formation.

We assume that H-bond disruption and formation involve respective energy release and absorption, such that each H-bond would create a small energy barrier in the conformation energy landscape. Based on this assumption, we constructed an energy landscape model of α_V_β_3_ with 7 energy barriers serially distributed between the bent and extended states, thereby creating 8 conformational states (one bent, 6 intermediate and one extended) (Fig. 6F). For the sake of simplicity, we further assumed that all energy barriers between the intermediate states were identical in shape and evenly distributed between the bent and extended conformations, hence having identical transition rates between any two adjacent intermediate states: *k*_–_ and *k*_+_. The respective rates of transition from the bent or extended state to their adjacent intermediate states were designated as 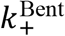 and 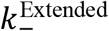 respectively. In our cell-free system, the only energy source that drives the conformational transitions is that microscopic thermal agitations from the macroscopically thermodynamically equilibrated environment. By assuming that the integrin structure does not provide any mechanism to regulate the directional tendency of conformational changes, integrins that reside in any intermediate state could transition bidirectionally towards either bending or unbending regardless of the previous direction of its immediate past transition.

By treating the stochastic conformational changes as a Markov process in a finite state space, including bent, intermediate, and extended states, we built a master equation 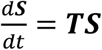, where ***S*** is the state occupancy vector. ***T*** is a [N+2]-by-[N+2] matrix of transition rates, in which N = 6 is the number of intermediate states. By solving the master equation, we obtained average time-to-transition <*t*_0±_> and switch time <*t*_sw±_> below (see Supp. Methods for details):

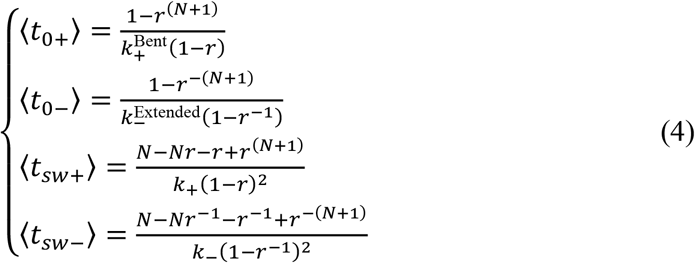

where 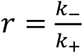. Assuming that the transition between adjacent states follows the Bell model^23^, all transition rates in Eq. 4 are regulated by force:

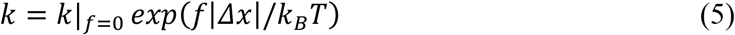

where *k*|_*f*=0_ is the value of *k* under zero force. |Δ*x*| and 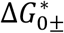 are, respectively, the distance and energy difference from the bottom of the energy well of any intermediate state to the top of its adjacent energy barrier in the energy landscape that takes the positive sign for unbending and the negative sign for bending.

Remarkably, our model fitted the experimental <*t*_sw±_> and <*t*_0±_> vs. force data measured in both cation conditions quite well (Fig. 5E,G,H), returning two sets of best-fit parameters, one for each cation condition. This allowed us to evaluate the parameters of the energy landscape, including differences between neighboring states: 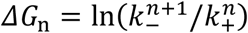(*n* = 0 (Bent), 1, 2, …N) (‘N+1’ represents ‘Extended’ state), and plot the energy landscape of α_V_β_3_ conformational changes (Supp. Table 2, Fig. 6F).

To further validate our model, we used Monte Carlo simulations to perform ‘mock runs’ based on this energy landscape. Gratifyingly, we were able to recreate integrin spontaneous bending and unbending conformational changes over time (Supp. Fig. 2A,B). Our multi-state model predicts that integrin may jump back-and-forth between adjacent states. Indeed, we observed that integrins occasionally paused in the middle of a bending process and reversed course to unbend in both Monte Carlo simulations (Supp. Fig. 2A) and BFP experiment (Supp. Fig. 2C). These results validate our energy landscape of α_V_β_3_ conformational changes.

## Discussion

Force-modulated bending and unbending conformational changes have previously been observed on cell surface integrins^6,7^. Here, we provide the first set of real-time single-molecule experimental data to show that purified integrin ectodomains are capable of undergoing force-modulated bending and unbending independent of cellular environment. Our results reveal very different biophysical characteristics for the two focal adhesion integrins α_5_β_1_ and α_V_β_3_. In the absence of tensile force, the α_5_β_1_ protein is nearly mono-stable with a probability of assuming the bent confirmation 3 orders of magnitude (*e*^7.2^) higher than that of assuming the extended confirmation in Ca^2+^. In the presence of force, however, α_5_β_1_ is rapidly switched to the extended conformation depending on whether the force is below or above a threshold of 7.4 pN where α_5_β_1_ becomes a bi-stable structure with 50-50 chance of being in either conformation. By comparison, even in the absence of force, the α_V_β_3_ protein is already bi-stable with a probability of being in the bent confirmation less than 3 times (*e*^0.99^) higher than that of being in the extended confirmation in Ca^2+^/Mg^2+^. The conformational changes of α_V_β_3_ are more gradually modulated by force in an “analogous” fashion as opposed to the “digital” fashion seen in the α_5_β_1_ case. Whereas bending and unbending of α_5_β_1_ can be modeled by a commonly used energy landscape of two wells separated by a barrier, which is simply ascribed to the Dudko-125 Hummer-Szabo model^29^, bending and unbending of α_V_β_3_ require a multi-state energy landscape to describe three orders of magnitude slower kinetics and lower force sensitivity. This newly developed model allows α_V_β_3_ to spontaneously bend and unbend over multiple steps, driven by the stochastic formation and disruption of a set of spatially distributed H-bonds in a reversible random sequence. The sequential formation and disruption of H-bonds serves as a seven-step ‘stair’ for α_V_β_3_ to temporarily ‘rest’ as it moves up- and down-stairs, thereby reducing the energy differential required in each step to the scale that can be provided by microscopic energy from thermal agitations despite that the macroscopic environment is in thermodynamic equilibrium. This model also provides a reasonable explanation for the long switching time *t*_sw±_ of α_V_β_3_, which is essentially the add-up of the time for transitioning across intermediate states. In the development of the α_V_β_3_ model, we only analyzed H-bonds for the sake of simplicity; however, other types of noncovalent interactions do exist to potentially mediate the integrin conformations, such as salt bridge, hydrophobic interaction, and thiol-disulfide exchange, which may be included in future studies. On the other hand, we are unable to investigate the structural basis of the force-modulated conformational changes of α_5_β_1_ due to the lack of its crystal structure. It is possible that noncovalent interactions at the atomic level involved in α_5_β_1_ conformational changes may either be much less in number, or be spatially synchronized and cooperative to allow concurrent disruption or formation, thereby composing a single large energy barrier.

Interestingly, the distinctive mechano-sensitivities of α_5_β_1_ and α_V_β_3_ may be evolved to support their respective biological roles in focal adhesion. Extending the prevailing view of integrin mechanosensing that requires the cooperation of multiple integrins and scaffold molecules (e.g., talin in the “molecular clutch” model^30^) to fulfill signal transduction, we speculate that integrin unbending may provide an additional mechanism for transducing outside-in mechano-signals, wherein the conformational change propagates intracellularly to induce integrin tailpiece separation and/or the association of cytoplasmic proteins^9^. The digital unbending of α_5_β_1_ by force allows the cell to quickly sense extracellular stretching above a threshold, and initiate α_5_β_1_ recruitment and clustering to form strong adhesion^3^. Furthermore, around the threshold force (7.4 pN), integrin α_5_β_1_ quickly switches back-and-forth between the bent and extended conformations (>10 Hz). Such high-frequency conformational changes could trigger fast oscillation in binding force magnitude, and therefore trigger the strong CMR effect^18^ of α_5_β_1_ to reinforce adhesion. On the other hand, with the analogous modulation of α_V_β_3_ (un)bending by force, the equilibrium of α_V_β_3_ conformations would be gradually shifted over a wide range of forces. This enables each α_V_β_3_ molecule to act as a ‘ruler’ for the cell to ‘measure’ the magnitude of extracellular stretching force and matrix rigidity. In support of this view, both MD simulations^31^ and experiment measurements^32^ have demonstrated that tensile forces exerted onto the headpiece of β_3_ integrins would directly cause the destabilization of the α-β transmembrane association and the rearrangement of integrin cytoplasmic tail independent of talin.

Outside-in mechano-signaling is a critical function of integrins on cell surface, for which multiple mechanistic models have been proposed. It was believed that to induce integrin mechano-signaling either requires integrins to cluster, so as to trigger rearrangement of cytoskeletal structure^33,34^, or alternatively, requires prior inside-out signaling to unbend the integrin for activation and ligand binding (“switch-blade” model^35^), and/or to activate intracellular scaffold proteins for signal transduction (e.g., talin in “molecular clutch” model^30^). In contrast to these conventional models, our current work and previous findings^6,7^ showed that bent integrins can also bind to ligands and that integrin unbending can be solely modulated by mechanical force. These results endorse the feasibility of a mechanism of single integrin outside-in mechano-signaling that does not require prior inside-out signaling, wherein integrin unbending plus force pulling allosterically cause integrin α/β tailpiece separation and/or integrin cluster re-arrangement^9,36^ to initiate intracellular signals.

Many molecular systems have been shown the capability of force-modulated reversible conformational transitions. For instance, complexes of T cell receptor (TCR) and pre-TCR with ligands have be observed to hop between compact and extended states under 5-15 pN tensile forces^37^. The mechanosensitive domain of platelet glycoprotein Ibα has been observed to unfold by >10 pN forces and refold when force is <12 pN^38^. Force has also been observed to regulate the unfolding and folding of protein L, pMHC, talin and titin^39-42^. Like integrins, multi-stability may be an intrinsic property of the structures of these molecules. Yet, α_V_β_3_ appears to be the only molecule identified so far that is capable of slow-kinetic sizable spontaneous conformational changes under a wide range of force without any energy source such as ATP, and hence requires a multi-state energy landscape model. Nonetheless, we cannot help to speculate that more molecules with a similar attribute should exist and await to be discovered. It will be interesting in the future to study these mechano-sensitive structures, not only to determine the exact shape of the intermediate states of their energy landscapes on the route of conformational changes, but also how they can accumulate and convert small-scale thermal energy into the work required for large-scale molecular conformational changes against force. Thus, the present work has not only provided actual examples but also critical concepts and useful design principles for engineering of protein biomechanical machines in the field of bionanotechnology^43^.

## Methods

### Proteins, antibodies and reagents

Previously described^15,30^ recombinant α_5_β_1_-Fc and trα_5_β_1_-Fc were generous gifts of Martin J. Humphries (University of Manchester, UK)^44^, α_V_β_3_-Hexa-His was a kind gift of Junichi Takagi (Osaka University, Japan)^24^, and FN biotinylated at the N-terminus was a kind gift of Andres J. Garcia (Georgia Institute of Technology, USA)^45^. The anti-FN mAb (HFN7.1) was from Developmental Studies Hybridoma Bank (Iowa City, IA). The anti-human Fc capturing mAb (GG-7) was from Sigma-Aldrich (St. Louis, MO). LIBS-2 was purchased from EMD Millipore (Billerica, MA). Anti-Penta-His antibody was purchased from Qiagen (Germany).

MAL-PEG3500-NHS and Biotin-PEG3500-NHS were from JenKem (Plano, TX). Nystatin, streptavidin-maleimide and BSA were from Sigma-Aldrich. Borosilicate glass beads were from Distrilab Particle Technology (RC Leusden, The Netherlands).

### AFM setup, preparation and experiment

Our AFM was built and calibrated in-house^15^. A petri-dish was directly mounted onto a piezo (P-363, Physik Instrumente, Karlsrube Germany), which was controlled by a computer program (Labview, National Instruments) with sub-nanometer spatial resolution through capacitive sensor feedback. A laser (Oz Optics, Ontario, Canada) was focused on the back of the cantilever (TM microscopes, Sunnyvale, CA) end, and deflected onto a photodiode (Hamamatsu, Bridgewater, NJ) to allow the cantilever deflection to be converted to force based on the cantilever spring constant^46^. To engage the integrins with FN, cantilever tips were incubated with 10-20 μg/ml FN overnight at 4 °C, rinsed, and incubated in Tris-buffered saline (50 mM Tris-Cl, 150 mM NaCl, pH 7.5) containing 1% bovine serine albumin (BSA) for 15 min at room temperature to block nonspecific binding^15^. For integrin coating, anti-Penta-His antibody was adsorbed on the petri-dish and then incubated with 10 μg/ml α_V_β_3_-Hexa-His, or GG-7 was adsorbed on the petri-dish by overnight incubation at 4 °C, rinsed, and incubated with 10 μg/ml α_V_β_3_-Hexa-His, α_5_β_1_-Fc or trα_5_β_1_-Fc for 30 min.

Some of the AFM experiment procedures have been described previously^15,18^. Briefly, the petri-dish was added with buffer of the desired cation composition. The piezo brought the petri-dish to contact the cantilever tip, retracted slightly and held the petri-dish close to the tip for 0.5 s to allow bond formation, and then retracted it at a speed of 200 nm/s. The presence of an adhesion event was reflected by a positive force signal in the force-time curves. The coating of the petri-dish was titrated to keep adhesion infrequent (<20%), a necessary condition for most of the adhesion events (>89%) to be mediated by single bonds^14^. For force-induced unbending and rebending measurements, the petri-dish was driven at a constant speed (200 nm/s) to load the bond to ∼20 pN and retract at the same speed to unload the bond. The (un)bending events were identified and parameters measured from the force-time traces (c.f., Fig. 1B). For CMR measurements, the petri-dish was driven to move cyclically so the integrin– FN bond underwent force loading and unloading and then held at a pre-set force (c.f., Fig. 4A,C)^18^. Lifetime was measured from the instant when the force reached the desired level to the instant of bond dissociation. The collected lifetime data were categorized into bins of successive force ranges, and averaged within each force bin to plot the lifetime curve. For force-ramp after a cyclic loading-unloading cycle with a high peak force, the piezo was retracted at a very low speed (1 nm/s) to allow observation of repetitive unbending and bending events over a prolonged period until bond rupture.

### RBC and glass bead preparation

Human blood (8–10 μl) was obtained from finger prick following a protocol approved by the Institutional Review Board of Georgia Institute of Technology (protocol number H12354), and RBCs were isolated and biotinylated by incubating with Biotin-PEG3500-NHS solution^6^. The biotinylated RBCs were then incubated with nystatin, which would swell the RBCs to near spherical shapes.

The procedure for bead functionalization has been described^47^. Briefly, after thiolation, glass beads were incubated with streptavidin-maleimide, anti-Penta-His antibody crosslinked with MAL-PEG3500-NHS, or LIBS-2 crosslinked with MAL-PEG3500-NHS overnight. Streptavidinylated beads were incubated with biotinylated FN solution for 2 h. Anti-Penta-His antibody coated beads were incubated with α_V_β_3_-Hexa-His solution for 3 h. LIBS-2 coated beads were used without further incubation. All beads after incubation were washed with and resuspended in phosphate buffer (27.6 g/L NaH_2_PO_4_·H_2_O, 28.4 g/L Na_2_HPO_4_).

### Platelet isolation

Procedures for collecting blood as approved by the Institutional Review Board of the Georgia Institute of Technology (protocol number H12354). Blood was collected from healthy volunteers into tubes containing anti-coagulant and activation-suppressing agents, and centrifuged at 200 g for 15 min to isolate platelet-rich plasma, which was centrifuged at 900 g for another 10 min to isolate the platelet pellet. The platelet pellet was resuspended in platelet washing buffer (4.3 mM K_2_HPO_4_, 4.3 mM Na_2_HPO_4_, 24.3 mM NaH_2_PO_4_, 113 mM NaCl, 5.5 mM D-Glucose, 10 mM theophylline, 20 U/mL Clexane, 0.01 U/mL apyrase, 1% BSA, pH 6.5), rested for 15 min and centrifuged again. Finally, the platelet pellet was resuspended into Hepes-Tyrode buffer (134 mM NaCl, 12 mM NaHCO_3_, 2.9 mM KCl, 0.34 mM sodium phosphate monobasic, 5 mM HEPES, and 5 mM glucose, 0.02 U/mL apyrase, 1% BSA, pH 7.4).

### BFP setup, preparation and experiment

Our BFP apparatus has been described previously^7,47^. A chamber mounted on an inverted microscope (Nikon TiE, Nikon) was filled with experimental buffer supplemented with 1% BSA to block nonspecific binding and cations (1 mM Ca^2+^/Mg^2+^ or 2 mM Mn^2+^). A biotinylated RBC was aspirated by a micropipette to act as a force transducer (Figs. 2A and 3E, *left*), the spring constant of which was set to 0.3 or 0.25 pN/nm^6^. A probe bead bearing FN or LIBS-2 was attached to the apex of the RBC via streptavidin-biotin interaction. An α_V_β_3_-functionalized bead or a platelet was aspirated by an opposing micropipette (Figs. 2A and 3E, *right*) as the target, and driven by a piezoelectric translator (Physical Instrument) to repeatedly touch with the probe bead and retract. The probe bead’s position was tracked by a high-speed camera.

The BFP measurement procedures for bond lifetime, (un)bending, and CMR are similar to AFM experiments. Briefly, a tensile force signal indicated an adhesion event between the probe bead and the target. FN coating on the probe bead was titrated to maintain infrequent adhesion (<20%)^14^. Upon the detection of an adhesion event, the target pipette was held at a desired position (reflected by the initial clamping force) to wait for the bond to dissociate.

### Molecular stiffness measurement

As previously described^6,7^, force vs. time data from AFM and BFP experiments were transformed to “force vs. extension” data (c.f., from Fig. 1B to 1C). The tensile force portion of the “force vs. extension” data was fitted by a line and the slope was taken as the stiffness of the integrin-FN complex. The value mainly reflects the integrin stiffness as the contribution from FN is negligible.

### Steered molecular dynamics (SMD) simulation of integrin α_V_β_3_

The ectodomain crystal structure of α_V_β_3_ (pdb id: 3ije)^48^ was used to perform the SMD simulation with GROMACS^49^. TIP3P model was used to depict water molecules. Na^+^ and Cl^-^ were added to neutralize the system and maintain the physiological salt condition (150 mM). The CHARM36 force field^50^ was used to describe the interactions of the protein and the solvent. CHARMM Additive All-Atom Force Field^51^ were used to describe the sugar. Simulations began by first minimizing the energy of the protein using steep decent methods, and then the system temperature was raised from 3 K to 300 K in an annealing simulation with controlled volume within 500 ps, following by another 500 ps simulation in NVT ensemble. Afterwards, we performed 1 ns simulation in NPT ensemble at 300 K and 1 atm. The temperature and pressure were controlled by a V-rescale thermostat and Parrinello−Rahman barostat, respectively^52^. In the annealing, NVT and NPT simulations, the positions of the heavy atoms of the integrin were restrained.

In the pulling simulation, the C-terminal Cα atom of β tail was restrained, and a group of atoms in the integrin head (Cα of residues 113-117, 151-156, 190-197, 244-250, 306-310, and 329-332 in βA domain of β subunit) was pulled at a speed of 0.1 nm/ns. Five independent pulling simulations were performed. The head-to-tail distance was minimally ∼3 nm and maximally ∼18 nm, consistent with the displacement of ∼13 nm observed in the BFP experiments.

In the position clamped simulation, C-terminal Cα atom of β tail was again restrained, and the clamping force was applied to the same group of atoms, but the pulling speed was set to 0. The bent (∼3 nm), extended (∼18 nm) and three partially extended (∼6, 11, and 16 nm) integrin structures with different head-to-tail distances were extracted from above simulation trajectories to monitor the number of H-bonds in a static molecular structure.

### Statistical Analysis

Statistical significance was assessed by unpaired or paired, two-tailed Student’s t-test or one-way ANOVA.

## Acknowledgements

We thank Andres J. Garcia (Georgia Institute of Technology, USA) for providing FN, Martin J. Humphries (University of Manchester, UK) for providing recombinant α_5_β_1_-Fc and trα_5_β_1_-Fc, and Junichi Takagi (Osaka University, Japan) for providing recombinant α_V_β_3_-Hexa-His. We thank Wilbur Lam (Georgia Institute of Technology, USA) lab for blood collection. This work was supported by National Institutes of Health grants R00HL153678 (Y.C.) and R01HL132019 (C.Z.), Army Research Office DOD W911NF-16-1-0257 (C.Z.) and a startup fund from The University of Texas Medical Branch. L.A.J. is an Australian Research Council DECRA Fellow (DE190100609) and a former National Heart Foundation of Australia postdoctoral fellow (101798).

## Author contributions

Y.C. supervised the study, designed and performed BFP experiments, analyzed data, helped develop the α_V_β_3_ model and wrote the paper; F.K. designed and performed AFM experiments, analyzed data, developed the α_5_β_1_ model and wrote the paper; Z.L. performed simulations, developed the α_V_β_3_ model and wrote the paper; L.A.J. provided critical suggestions and wrote the paper; C.Z. supervised the study, designed experiments and wrote the paper. Research activities related to this work were complied with relevant ethical regulations.

