## Supplementary information for "Force-regulated spontaneous conformational changes of integrins α_5_β_1_ and α_V_β_3_"

**Supplementary Figures**

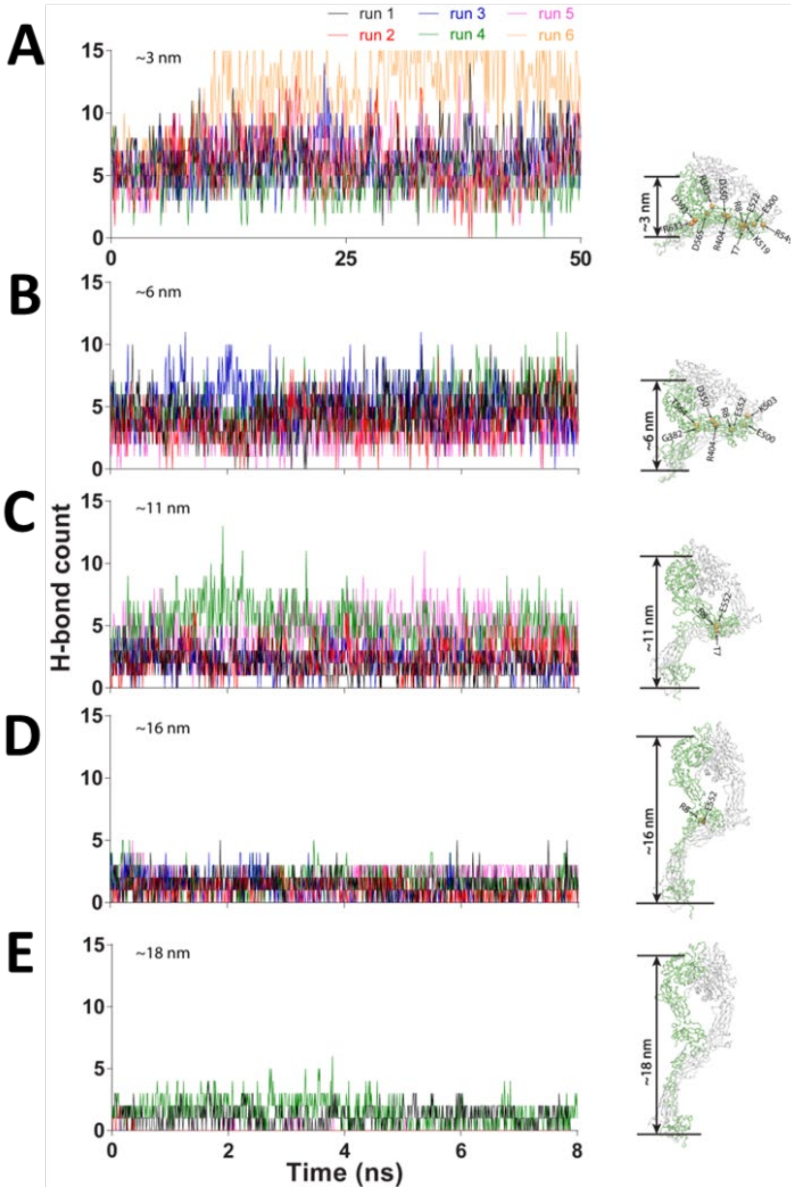


**Supplementary figure 1. Using MD simulation to track the number of H-bonds between αvβ3 headpiece and tailpiece over time.** The integrin head-to-tail distance was fixed respectively at ~3 (A), ~6 (B), ~11 (C), ~16 (D) and ~18 (E) nm (*right*). Each panel exhibit the traces of 6 repeated simulations, marked by different colors.

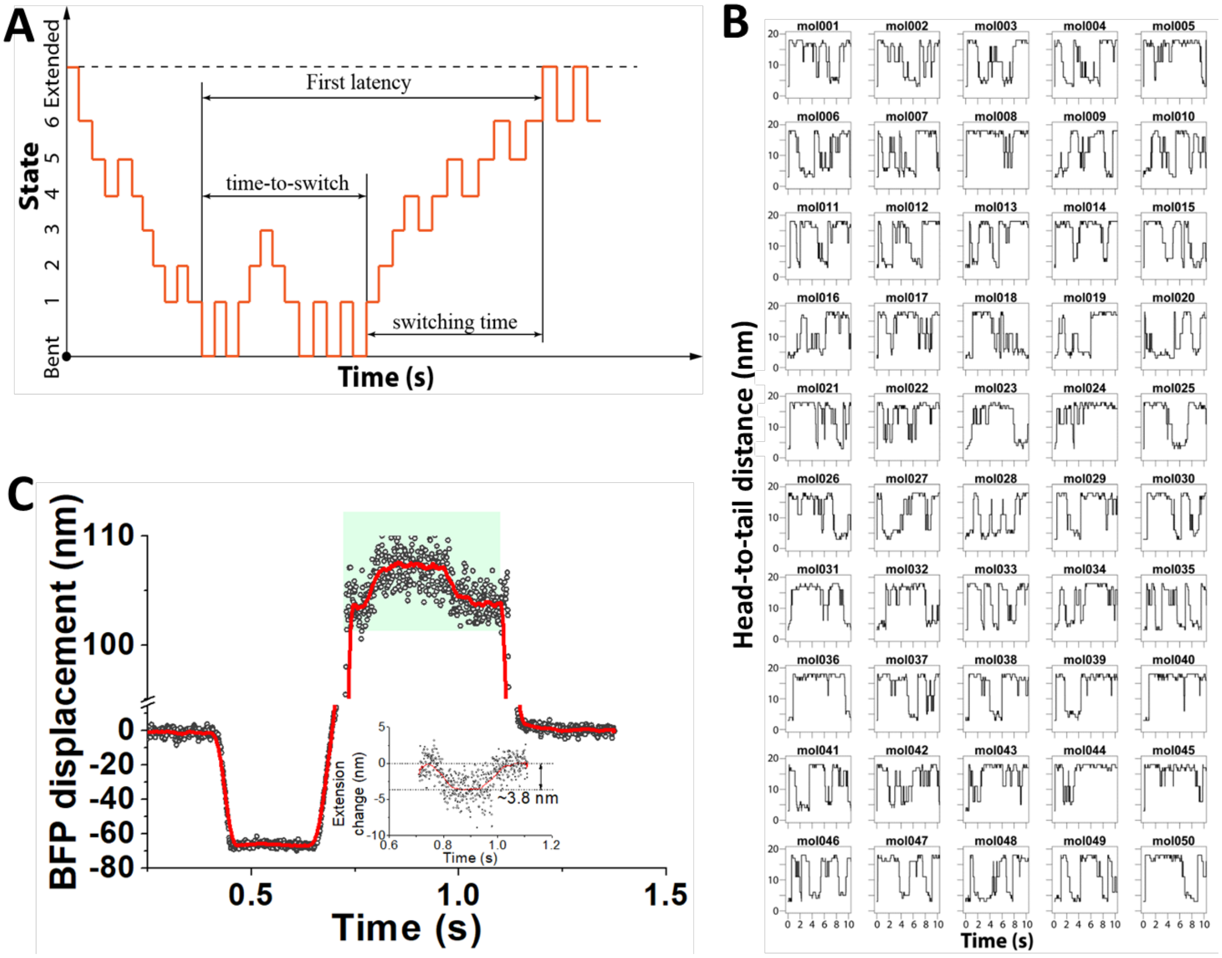


**Supplementary figure 2. Simulation demonstration and experimental evidence of multi-state model of $\alpha_V\beta_3$ conformational changes.** A. A representative 'mock run' of $\alpha_V\beta_3$ conformational dynamics using Monte Carlo (MC) simulation, depicting an integrin fulfilling a bending event followed by an unbending event. Definition of time-to-switch and switching time of the unbending process were annotated on the graph. B. Integrin head-to-tail distance vs. time in 50 runs of MC simulation. Each panel represents one run. The head-to-tail distance was obtained based on the integrin's conformational state. C. A representative BFP force *vs*. time signal showing a partial and unfinished unbending event interrupted by an ensuing bending back.

**Supplementary Tables**

| Probe | Condition | #Lifetime events | Unbending #Events | Bending #Events |
|---|---|---|---|---|
| FN | $Mn^{2+}$ | 1424 | 185 | 43 |
| | $Ca^{2+}/Mg^{2+}$ | 1847 | 164 | 70 |
| | $Ca^{2+}/Mg^{2+}$/ LIBS-2 | 183 | 5 | 0 |
| LIBS-2 | $Ca^{2+}/Mg^{2+}$ | 186 | 0 | 0 |

**Supplementary table 1. Statistics of BFP-observed $\alpha_V\beta_3$ lifetimes and unbending and bending events while binding to FN- and LIBS-2-coated probe beads.**

| | $k_{Bent}$ | $\Delta x_{Bent}$ | $k_{+}$ | $\Delta x_{+}$ | $k_{Extend}$ | $\Delta x_{Extend}$ | $k_{-}$ | $\Delta x_{-}$ |
|---|---|---|---|---|---|---|---|---|
| $Ca^{2+}$/ $Mg^{2+}$ | 2.13E+01 | ~0 | 4.84E+01 | 9.27E-02 | 1.93E+00 | 1.31E-01 | 3.83E+00 | ~0 |
| $Mn^{2+}$ | 6.95E+00 | ~0 | 2.38E+01 | 1.01E-01 | 2.56e+00 | 3.57E-02 | 2.16E+01 | ~0 |

**Supplementary table 2. Fitted kinetics and energy landscape parameters of $\alpha_V\beta_3$ bending and unbending.**

**Supplementary Methods**

*Solution of the multiple state transition kinetics*

Since the transition of integrins between different states (bent, intermediate and extended) are manipulated by stochastic thermodynamic energy, an integrin in any given intermediate state could transition towards both bending and unbending. We argue that the long switching time $t_{sw\pm}$ is essentially the add-up of the time-to-switch for all the jumps among intermediate states. By treating the stochastic conformational change as a Markov process in a finite state space, including bent, intermediate, and extended states. The dynamics of each state can be described as:

$$\frac{\mathrm{d}\mathbf{S}}{\mathrm{d}t} = \mathbf{TS},$$

where **S** is the state occupancy vector. **T** is a N+2-by-N+2 matrix of transition rates, where N is the number of the intermediate states.

In the occupancy vector, $S_{\mathrm{Bent}}$, $S_i$ ($i$=1,3…N), and $S_{\mathrm{Extended}}$ represent the bent, the $i^{\mathrm{th}}$ intermediate, and the extended conformations. The transition rate matrix is defined by the transition rates:

$$T = \begin{pmatrix} -k_+^{\mathrm{Bent}} & k_-^1 & 0 & 0 & \ldots & 0 \\ k_+^{\mathrm{Bent}} & -\left(k_-^1 + k_+^1\right) & k_-^2 & 0 & \ldots & 0 \\ 0 & k_+^1 & -\left(k_-^2 + k_+^2\right) & k_-^3 & \ldots & 0 \\ & & \ddots & & & \\ \ldots & \ldots & \ldots & k_+^{\mathrm{N}-1} & -\left(k_-^{\mathrm{N}} + k_+^{\mathrm{N}}\right) & k_-^{\mathrm{Extended}} \\ 0 & \ldots & \ldots & 0 & k_+^{\mathrm{N}} & -k_-^{\mathrm{Extended}} \end{pmatrix}$$

(4)

$k_+^{\mathrm{Bent}}$, $k_-^{\mathrm{Extended}}$ are the rates of escaping from the bent or extended states to the nearby intermediate states. $k_+^i$, $k_-^i$ ($i$=1,2…N) are respectively the transition rate of the $i$th intermediate state (indicated by the superscript) along the transition pathway. Marks "+" and "–" respectively represent the transition direction toward unbending and bending. All the other entries in **T** equal to zero.

The mathematical definition of $t_{0\pm}$ and $t_{sw\pm}$ could then be given as follows. The sums ($t_{0\pm}+t_{sw\pm}$) of the time-to-switch $t_{0\pm}$ and switching time $t_{sw\pm}$ are the first latency to complete a full transition between two stable end-states (from bent to extended or *vice versa*; Supp. Fig. 5A). Notably, before the fulfillment of a complete transition, the molecule is allowed to undergo multiple incomplete transitions ending by returning to the initial end-state. The $t_{0\pm}$ covers all the time consumed by these back-and-forth incomplete transitions, and finally when a complete transition starts, covers the latency for the end-state integrin to enter the first intermediate state (Supp. Fig. 5A). On the other hand, $t_{sw\pm}$ covers the rest of the time consumed by the molecule to complete the full transition, in which the molecule passes through all the intermediate states (mostly likely back-and-forth) and eventually arrives in the other end-state (Supp. Fig. 5A). By definition, a full transition requires the molecule to transit from one end-state to the other, during which the molecule cannot return to the initial end-state. Therefore, we argue that the first latency to complete transitioning from the intermediate state adjacent to end-state A to end-state B without the existing of end-state A is equal to the switching time $t_{sw\pm}$, which provides a simpler way to calculate $t_{sw\pm}$. In this case, the transition matrix is the submatrix of the original one, either deleting the first column and row or deleting the last column and row from the original matrix. The first latency to complete a full transition has been solved previously[1], where the average of sums ($<t_{0\pm}+t_{sw\pm}>$) and the switching time ($<t_{sw\pm}>$) can be written as:

$$\begin{cases} \left\langle t_{0+} + t_{sw+} \right\rangle = \sum_{i=1}^{N} \dfrac{1+\sum_{j=i}^{N}\prod_{k=i}^{j} r_k}{k_+^{i-1}} + \dfrac{1}{k_+^{N}} \\ \left\langle t_{0-} + t_{sw-} \right\rangle = \sum_{i=1}^{N} \dfrac{1+\sum_{j=1}^{i}\prod_{k=j}^{i} \frac{1}{r_k}}{k_+^{i+1}} + \dfrac{1}{k_-^{1}} \\ \left\langle t_{sw+} \right\rangle = \sum_{i=2}^{N} \dfrac{1+\sum_{j=i}^{N}\prod_{k=i}^{j} r_k}{k_+^{i-1}} + \dfrac{1}{k_+^{N}} \\ \left\langle t_{sw-} \right\rangle = \sum_{i=1}^{N-1} \dfrac{1+\sum_{j=1}^{i}\prod_{k=j}^{i} \frac{1}{r_k}}{k_+^{i+1}} + \dfrac{1}{k_-^{1}} \end{cases} \tag{5}$$

where $r_i = \dfrac{k_-^i}{k_+^i}$, $k_+^0 = k_+^{\text{Bent}}$, and $k_-^{N+1} = k_-^{\text{Extended}}$

By subtracting the switching time from the first latency to complete a full transition, one can obtain the time-to-switch. Thus, the time-to-switch and switching time can be summarized as below:

$$\begin{cases} \left\langle t_{0+} \right\rangle = \dfrac{1+\sum\limits_{j=1}^{\mathrm{N}} \prod\limits_{k=1}^{j} r_k}{k_+^{\mathrm{Bent}}} \\ \left\langle t_{0-} \right\rangle = \dfrac{1+\sum\limits_{j=1}^{N} \prod\limits_{k=j}^{N} \dfrac{1}{r_k}}{k_+^{\mathrm{Extended}}} \\ \left\langle t_{sw+} \right\rangle = \sum\limits_{i=2}^{\mathrm{N}} \dfrac{1+\sum\limits_{j=i}^{\mathrm{N}} \prod\limits_{k=i}^{j} r_k}{k_+^{i\text{-}1}} + \dfrac{1}{k_+^{\mathrm{N}}} \\ \left\langle t_{sw-} \right\rangle = \sum\limits_{i=1}^{\mathrm{N}\text{-}1} \dfrac{1+\sum\limits_{j=1}^{i} \prod\limits_{k=j}^{i} \dfrac{1}{r_k}}{k_+^{i+1}} + \dfrac{1}{k_-^{1}} \end{cases} \tag{6}$$

Each energy barrier on the transition pathway corresponded to the breakage/formation of H-bond. To simplify our model, we assumed all the energy barriers among the intermediate states to be identical. Therefore, the transition rates $k_+^i$, $k_-^i$ ($i$=1…N) were identical between every two neighboring states, and were labeled as $k_-$ and $k_+$, respectively. In turn, the ratio of transition rates, $r_i = \dfrac{k_-^i}{k_+^i}$, were also identical and were labeled as $r$. With these simplifications, we re-wrote the model as Eq. (3).
